# VIP interneurons in mouse whisker S1 exhibit sensory and action-related signals during goal-directed behavior

**DOI:** 10.1101/2021.10.08.463283

**Authors:** Deepa L. Ramamurthy, Andrew Chen, Patrick C. Huang, Priyanka Bharghavan, Gayathri Krishna, Kayla Casale, Daniel E. Feldman

## Abstract

Vasoactive intestinal peptide-expressing (VIP) interneurons, which constitute 10-15% of the cortical inhibitory neuron population^1,2^, have emerged as an important cell type for regulating excitatory cell activity based on behavioral state. VIP cells in sensory cortex are potently engaged by neuromodulatory and motor inputs during active exploratory behaviors like locomotion and whisking, which in turn promote pyramidal cell firing via disinhibition^3-5^. Such state-dependent modulation of activity by VIP cells in sensory cortex has been studied widely in recent years. However, the function of VIP cells during goal-directed behavior is less well understood. It is not clear how task-related events like sensory stimuli, motor actions, or reward activate VIP cells in sensory cortex since there is often temporal overlap in the occurrence of these events. We developed a Go/NoGo whisker touch detection task which incorporates a post-stimulus delay period to separate sensory-driven activity from action- or reward-related activity during behavior. We used 2-photon calcium imaging to measure task-related signals of L2/3 VIP neurons in S1 of behaving mice. We report for the first time that VIP cells in mouse whisker S1 are activated by both whisker stimuli and goal-directed licking. Whisker- and lick-related signals were spatially organized in relation to anatomical columns in S1. Sensory responses of VIP cells were tuned to specific whiskers, whether or not they also displayed lick-related activity.

## RESULTS AND DISCUSSION

Head-fixed VIP-Cre; Ai162D (TIGRE2.0-GCaMP6s) mice with 9 whiskers inserted into a piezo actuator array were trained to lick in response to deflection of a randomly chosen whisker (Go trials) to receive a reward, and to suppress licking on trials in which no whisker was deflected (NoGo trials, which were neither rewarded nor punished) (**Figure 1A-B**). Mice readily learned to perform this task, in an average of 20.3 ± 1.19 days (**Figure 1C**) following initial acclimation. Use of a delay period successfully shifted onset of licks in the trial beyond the window for analysis of stimulus-evoked responses (median first lick time = 1.52 ± 0.19 s), with only 12.8 ± 2.02% trials per session aborted due to early licks (**Figure 1D-E**).

**Figure 1.**
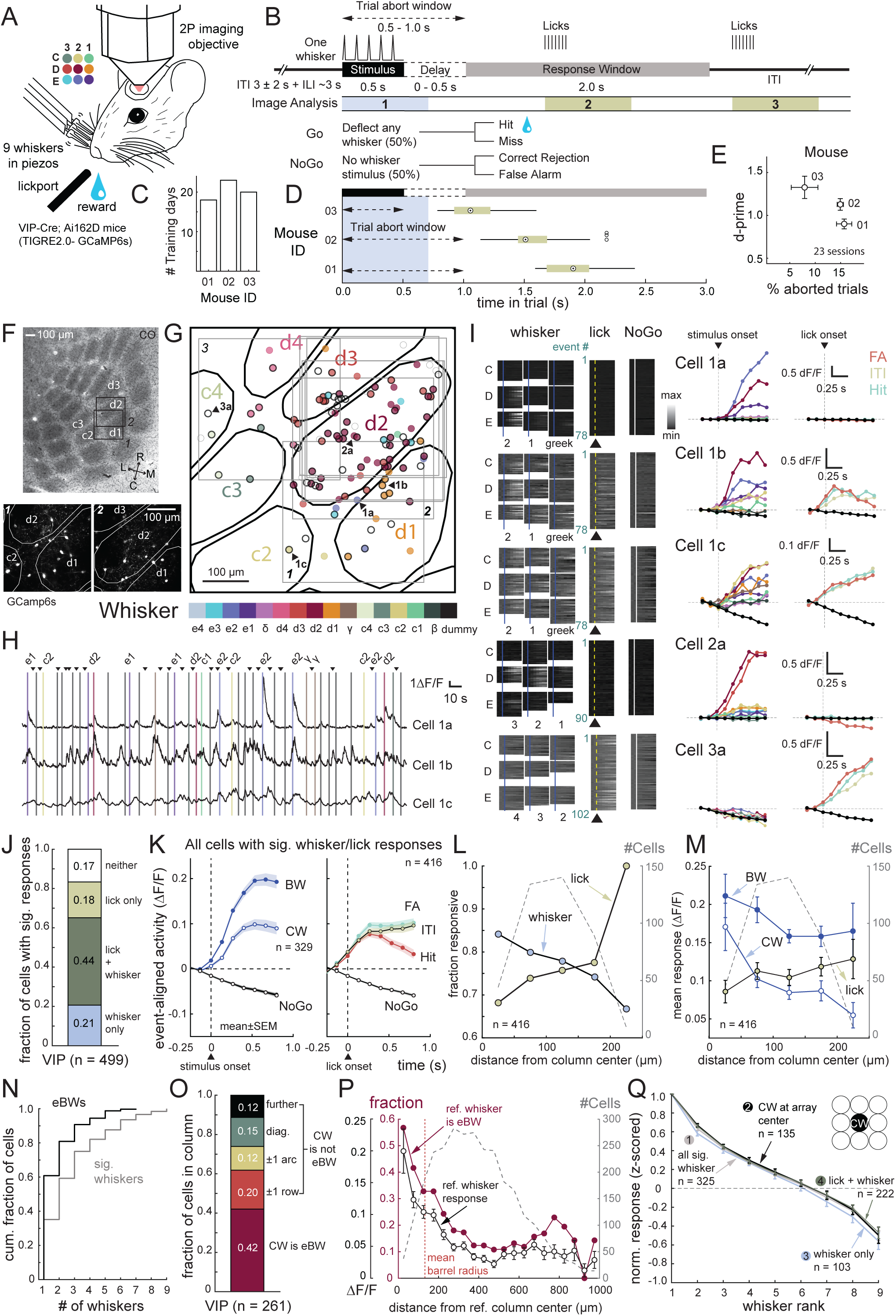
Sensory and action-related responses of VIP interneurons in mouse whisker S1 during goal-directed behavior. **A**. Head-fixed mice had 9 whiskers inserted in a piezo actuator array and were rewarded for licking when they detected any single-whisker deflection. **B**. Behavioral trial structure. On each trial, one randomly chosen whisker was deflected (Go trials) or no whisker was deflected (NoGo trials). Mice were rewarded for licking in the response window on Go trials (hits) but were neither rewarded nor punished for licking on NoGo trials (false alarms). The identity of the deflected whisker on Go trials was randomly selected. A delay period between stimulus and response window served to separate sensory-related from action- or reward-related neural signals. Windows for image analysis was aligned to (1) stimulus onset, (2) first lick of the lick bout in the response window and (3) first lick of lick bouts in the intertrial interval. **C**. Number of sessions (after initial rig acclimation) to reach expert performance. Imaging was commenced after behavioral performance was maintained at d’ ≥ 1 for 3 consecutive sessions. **D**. Median time of first licks across all imaging sessions. **E**. Mean d-prime across all imaging sessions. **F**. Two L2/3 imaging fields from a VIP-Cre; Ai162D (TIGRE 2.0-GCamp6s) mouse overlaid on a cytochrome oxidase-stained L4 section showing barrels (top). Bottom, column boundaries projected onto the two imaging fields. **G**. VIP cells (circles) from 7 imaging fields in one mouse. Whisker-responsive cells are in color (color denotes the best whisker). Lick-responsive cells have a black outline. The best whisker was identified as the CW if the CW was among the cell’s equivalent best whiskers (eBWs). **H**. Example ΔF/F traces from 3 VIP cells that were imaged together in one field. Bars, whisker deflections. Triangles, licks. **I**. Whisker receptive fields and lick-related activity for 5 example VIP cells, including the 3 cells shown in **(H)**. Left, ΔF/F traces for each trial for whisker stimuli, lick events, and dummy piezo stimuli. Middle, median ΔF/F trace for each whisker (color coded for whisker identity), and licks (color coded for FA, Hit or ITI licks). **J**. Proportion of L2/3 VIP cells that were whisker or lick responsive. **K**. Mean ΔF/F trace for all VIP cells with either significant whisker- or lick-related activity, aligned to whisker stimulus onset or lick onset. **L**. Proportion of VIP cells that were whisker- or lick-responsive as a function of cell distance from the CW column center. For septal cells, distance was measured from nearest column center. **M**. Same, but for mean response (ΔF/F) to whisker deflections and licks (baseline subtracted, see Methods). **N**. Receptive field size of VIP cells, quantified as number of significant whisker and eBWs. **O**. Distribution of BW identity for L2/3 VIP cells located within barrel columns (septal cells excluded). **P**. Proportion of cells tuned to a reference whisker, and mean response to that whisker, as a function of distance from the reference whisker column. **Q**. Mean rank-ordered receptive field for all whisker-responsive cells, calculated after ranking whiskers from strongest to weakest for each cell, with responses normalized to baseline activity.

We performed 2p imaging through a chronic cranial window during behavior in expert mice. Imaged cells were localized relative to the anatomical boundaries of barrel columns by post hoc histological staining and alignment with surface vasculature (**Figure 1F**). This allowed us to examine the map organization of task-related signals in VIP cells (**Figure 1G**). We measured VIP cell activity (ΔF/F) aligned to sensory stimuli (whisker deflections) and goal-directed actions (licks). Individual cells showed significant elevations in activity following whisker deflections, licks, or both. Previous studies showed whisker responses in VIP cells, but did not assess whisker tuning^6-9^. We observed prominent whisker somatotopic tuning for VIP cells, as well as lick responses (**Figure 1H-I**). At the population level, similar proportions of VIP cells were whisker-responsive (65.1%) and lick-responsive (62.7%), with most cells (44.5% of total) being activated by both, and fewer cells being either solely whisker-responsive (20.6%) or lick-responsive (18.2%) (**Figure 1J**). On average, both whisker stimuli and licks were strong drivers of VIP cell activity, although the strongest whisker responses had a higher magnitude than lick responses (**Figure 1K**). Lick-related responses (0.08 ± 0.01 ΔF/F) were comparable in magnitude to the mean response to deflection of the columnar whisker (CW; 0.07 ± 0.01 ΔF/F; p = 0.292, permutation test for difference in means) and were ∼50% of the mean response to each cell’s best whisker (BW; 0.14 ± 0.01 ΔF/F; p = 0.01, permutation test for difference in means. See **Figure S1A** for dynamics of activity in ITI vs. trial period). Lick-related activation of VIP interneurons has not previously been reported in sensory cortex, although there is evidence for this in dorsomedial prefrontal cortex of mice^10^. Lick-related activity in our study was not simply due to activation of VIP cells by whisker movements that accompany licking, because in our task VIP responses to spontaneous whisking events in the absence of licking were weak and impacted a smaller fraction of VIP cells (**Figure S1B**). This prominence of sensory and lick responses over whisking-related responses may reflect the stationary nature of our passive whisker task^11^.

Reinforcement signals in VIP cells have previously been reported in sensory cortex^4^. Therefore, we tested whether reward delivery on Hit trials would elevate lick-related VIP cell activity over that seen on false alarm (FA) trials or during spontaneous licking during the intertrial interval (ITI), which lack reward. On average, reward did not excite VIP cells – lick-related activity was relatively suppressed rather than elevated on hit trials (0.05 ± 0.01 ΔF/F), in comparison to FA licks (0.09 ± 0.01 ΔF/F; p <0.001, permutation test for difference in means, FDR correction for multiple comparisons; **Figure 1K**) or ITI licks (0.08 ± 0.01 ΔF/F; p = 0.346, permutation test for difference in means, FDR correction for multiple comparisons). Thus, reinforcement did not activate VIP cells in S1 in the context of our whisker detection task.

We examined whether whisker touch vs. lick signals in VIP cells are spatially organized relative to column boundaries in S1. Whisker- and lick-responsiveness of VIP cells both showed a dependence on their location relative to the center of the nearest barrel column, with a 50-100 μm resolution (**Figure 1L-M**). Whisker-vs. lick-responsiveness were inversely related (r = −0.96, p = 0.01, Pearson’s correlation) – a greater fraction of whisker-responsive cells was found near the center of the barrel (center, 0-50 μm: 84.1% whisker vs. 66.7% lick), while lick-responsive cells were more common near barrel edges (edge, 150-250 μm: 70.4% whisker vs. 88.8% lick; **Fig. 1L**). The magnitude of whisker- and lick-evoked responses also showed a similar center-dominance of whisker-evoked responses (center, 0-50 μm: CW = 0.17 ± 0.03 ΔF/F vs. lick = 0.09 ± 0.02 ΔF/F; p = 0.012, permutation test for difference in means) and edge-dominance of lick-evoked responses (edge, 150-250 μm: CW = 0.08 ± 0.01 ΔF/F vs. lick = 0.12 ± 0.01 ΔF/F; p = 0.013, permutation test for difference in means**; Figure 1M**). This opposing spatial organization could arise from spatial segregation at the level of inputs or outputs of distinct pathways associated with lick vs whisker signals. Among known inputs to mouse whisker S1^12^, one possible source of lick-related signals could be basal forebrain cholinergic neurons, which are excited by licking^13^ and have been shown to activate cortical VIP cells through acetylcholine release^14^.

Whisker selectivity of VIP cell responses was previously unknown in S1. Whisker-responsive VIP cells showed somatotopic whisker tuning (**Figure 1I, K**). On average, a mean of 2.73 ± 0.11 whiskers (out of 9) drove significant responses in each VIP cell, and 1.75 ± 0.07 whiskers drove statistically equal responses that were greater than all the rest (equivalent best whiskers, EBW**; Figure 1N**). Thus, VIP cells were tuned to specific whiskers. In mouse barrel cortex, L2/3 pyramidal cells tuned to different whiskers are intermixed in each column in salt-and-pepper organization^15-16^. We examined the tuning organization of VIP cells in the whisker map (**Figure 1G**). 41.8% of VIP cells in a given column were tuned the CW, 20.3% were tuned to neighboring whiskers in the same row, 11.5% in same arc, 14.9% at diagonal positions and 11.5% at further on the whisker pad (**Figure 1O**). The fraction of VIP cells with tuning that matches the CW is similar to that for PYR cells in awake mice^16^. This fraction (% CW/BW match) declined as function of distance from the column center (**Figure 1P**). We also examined the spatial profile of VIP cell activity in S1 evoked by deflection of a single whisker. To do this, we quantified each cell’s response to a reference whisker, and then examined this response across cells as a function of cell location relative to the center of that whisker’s column (**Figure 1P**). This average whisker response was greatest at the reference column center, and gradually fell off across the width of multiple barrel columns. Thus, VIP cells showed whisker tuning with average correct somatotopy in S1, but with an intermixed organization with very high local scatter, similar to PYR cells in mouse S1^16^. The above analysis of whisker tuning included both VIP cells driven only by whisker stimuli and those activated by both whisker stimuli and licks. These two subpopulations, when considered separately, showed identical whisker receptive field structure quantified by rank-ordered whisker tuning curves, and identical BW tuning preference (BW tuning preference for purely whisker-responsive cells = 0.52 ± 0.02; for whisker- and lick-responsive cells = 0.48 ± 0.01; p = 0.092, Wilcoxon rank-sum test; **Figure 1Q**). Thus, cells that were responsive to both whisker stimuli and licks did not correspond to a distinct population of cells that were less selective in their responses overall. Our measure of tuning width of VIP cells was not affected by limited sampling of whiskers in the receptive field due to positioning of the CW at the edges of the piezo actuator array that delivers whisker stimuli. This is shown in the rank-ordered receptive field for only the whisker-responsive cells which had their CW at the center of the piezo array (BW tuning preference = 0.48 ± 0.01), which was not significantly different from the remaining dataset (BW tuning preference = 0.51 ± 0.02; p = 0.085, Wilcoxon rank-sum test). Whisker tuning of VIP cells (BW tuning preference = 0.49 ± 0.01; see **Figure S1C** for CW centered tuning curves) was slightly broader than previously reported for PYR cells in awake mice^16^, consistent with VIP tuning for other sensory features in visual^17^ and auditory cortex^18^.

Together, our results demonstrate activation of VIP cells in S1 primarily by two task-related events – sensory stimuli (whisker deflections) and goal-directed actions (licking) – during a whisker detection task that includes a delay period. Whisker tuning in VIP cells suggests a possible functional role in shaping whisker sensory coding in S1 beyond global state-dependent modulation, perhaps similar to recent studies of VIP cell contribution to contrast-dependent surround suppression in mouse visual cortex^19^. While lick-related activity in VIP cells has not previously been reported in S1, a number of studies have reported subpopulations of PYR cells in S1 acquiring lick-related responses following sensorimotor learning of whisker-based tasks that involve licking^20-22^. Our finding that VIP cells show lick-related activity raises the possibility of their involvement in sensorimotor learning or in processes that facilitate execution of goal-directed behavior.

## ACKNOWLEDGMENTS

We thank Katherine Smith, Stephanie Richards, Tasfia Rashid and Katie Lin for mouse colony management. This work was funded by NIH grant F32NS114327 (DLR), a UC Berkeley SURF L&S fellowship supported by the Anselm M&PS Fund (AC) and NIH grant R37 NS092367 (DEF).

## AUTHOR CONTRIBUTIONS

D.L.R. and D.E.F. designed the study. D.L.R., A.C., G.K., and K.C. performed the experiments. D.L.R., A.C., P.C.H, and P.B. analyzed the data. D.L.R. and D.E.F. wrote the manuscript.

## DECLARATION OF INTERESTS

The authors declare no competing interests.

## Notes

### Competing Interest Statement

The authors have declared no competing interest.

