## Supplementary Materials and Methods for "VIP interneurons in mouse whisker S1 exhibit sensory and action-related signals during goal-directed behavior"

#### Animals

All procedures were approved by the UC Berkeley Animal Care and Use Committee and were in accordance with NIH guidelines. Data included for VIP interneurons was obtained from 23 imaging experiments across three VIP-Cre; Ai162D (2 male, 1 female), generated by crossing VIP-Cre mice (JAX #10908) and Ai162D mice (TIGRE2.0-GCaMP6s, JAX #031562), obtained from The Jackson Laboratory. Supplemental analyses include data from pyramidal (PYR) cells collected across 4 experiments in one male Drd3-Cre; Ai162D mouse, generated by crossing Drd3-Cre mice (Gensat MMRRC: 034610) with the Ai162D line.

#### Surgery

Adult mice (2-3 months) were anesthetized with isoflurane (1.0-3.0%), with body temperature maintained at 37°C. Pre-operative dexamethasone (2 mg/kg, to reduce inflammation), meloxicam (5-10 mg/kg, for prophylactic analgesia) and enrofloxacin (10 mg/kg, to prevent infection) were administered, sterile technique was followed during the surgery. A lightweight (<3 g) head plate containing a 6 mm aperture was affixed to the skull using cyanoacrylate glue and Metabond (C&B Metabond, Parkell). Intrinsic signal optical imaging (ISOI) was used to localize either C-row (C1, C2, C3) or D-row (D1, D2, D3) barrel columns in S1[1], and a 3 mm craniotomy was made over either the C2 or D2 column, using a biopsy punch. The craniotomy was covered by placing a 3 mm diameter glass coverslip (#1 thickness, CS-3R, Warner Instruments) over the dura, and sealed with Metabond to form a chronic cranial window. Mice were allowed to recover on a heating pad and monitored until sternal and ambulatory. Buprenorphine (0.05 mg/kg) was given subcutaneously for post-operative analgesia, and mice were returned to the home cage. Mice were allowed to recover for 1 week before behavioral training commenced. After recovery, ISOI was repeated through the cranial window to verify the locations of barrel columns prior to the start of imaging experiments.

**Behavioral task**

To motivate behavior, mice were water regulated. The daily water ration (typically 0.8-1.5 mL) was calibrated for each mouse to achieve a stable target body weight of 85% of pre-training body weight. Mice were trained to perform a Go/NoGo whisker detection task. During the task, the mouse was head-fixed, with the body resting on spring-mounted stage[2]. Nine whiskers were inserted in a 3 x 3 piezo array (consisting of nine independent, calibrated piezoelectric actuators mounted with pipet tips to hold individual whiskers) such that the tips of the piezos were positioned ~5 mm from the face. A plastic shield prevented piezos from contacting other neighboring whiskers. The array was centered on a D-row or C-row whisker, and each whisker was held in place with a small amount of rubber cement applied at the start of each session. A lickport with a capacitive lick sensor was placed in front of the mouse, and used to deliver water rewards (4 µl, mean reward volume). Mice were briefly anesthetized with isoflurane (0.5-2.0%) at the start of each session for head-fixation and whisker insertion, and data collection began after all observable effects of anesthesia had fully subsided. Behavior was conducted in the dark, with 850 nm IR illumination for video monitoring. A masking noise designed to conceal sounds from the piezo actuators was played in the background throughout the task. Task control was performed using an Arduino Mega 2560 microcontroller board (Arduino), and custom routines in Igor Pro (WaveMetrics) were used for user input and task monitoring.

<u>Task structure</u>

On each trial, one randomly selected whisker was deflected (Go trials) or no whisker was deflected (NoGo trials). The number of Go trials and NoGo trials was balanced (50% each). The whisker stimulus on each Go trial consisted of a train of 5 deflections with a 100 ms interval between them. Each individual deflection consisted of a 300 μm amplitude (6º angular deflection) rostrocaudal ramp-and-return movement with 2 ms rise/fall time and 10 ms duration. On each NoGo trial, the same stimulus occurred on a dummy piezo that did not have any whisker attached to it. Trial onset was irregular – the inter-trial interval (ITI) was a 3 ± 2 s, after which mice were required to suppress licking below an inter-lick interval (ILI) threshold for ~ 3 s to initiate the next trial. Mice were rewarded for licking within the response window (2.0 s) on Go trials but were neither rewarded nor punished for licking on NoGo trials. Behavioral outcomes on each trial were recorded as hit, miss, false alarm or correct rejection.

To separate sensory-driven activity from action- or reward-related neural signals during goal-directed behavior, we adopted two task parameters which were effective to promote licking with a delay following stimulus onset. One was a trial abort window (0.5-1.0 second) which included the stimulus window (0.5 sec) with or without an additional post-stimulus delay period (0.0-0.5 s). Licking during the trial abort window caused termination of the trial. The second was a graded reward delivery paradigm to shift the onset of response licks – reward size on hit trials was larger the later mice lick in the response window (linear ramp in reward volume during the first 50-75% of the response window duration, which then plateaus to the maximum reward volume). Aborted trials were excluded from analyses.

Training stages

Behavior was shaped gradually, through a series of training stages (lasting 1-5 days each). In Stage 1, mice were acclimated to experimental rig and handling by trainers. In Stage 2, they were acclimated to head-fixation and licking at the port to obtain water rewards. Full reward volume was provided and reward delivery was cued by a blue light, until the final stages when the training of delayed licking was performed. In Stage 3, reward (cued by a blue light) was automatically delivered if mice suppressed licking below ILI threshold for ~3 seconds. In Stage 4, the whisker stimulus was introduced, with reward (cued by a blue light) automatically delivered in the response window regardless of the actions of the mouse. 50% NoGo trial were introduced. When mice began shifting licks such that they preceded reward delivery, they were advanced to the next stage. In Stage 5, the task shifted to operant mode (without delayed licking). Reward was provided only if the mouse licked in the response window. Learning progress was evaluated by the divergence of lick probability on Go and NoGo trials. In Stage 6, training for delayed licking was introduced. Trial abort window was incrementally lengthened, in tandem with gradual shifts in the reward plateau time within the response window. A limit was placed on the number of consecutive Go or NoGo trials to facilitate learning of delayed licking. Learning progress was evaluated measuring the median time of first licks in trials, relative to stimulus onset. In Stage 7, the final version of the whisker detection task was implemented, with fully randomized Go/NoGo trials and trial abort window and reward plateau parameters at their final values. Mice were considered achieve criterion performance when performance at this stage remained stable at d'>1 for three consecutive sessions. Fully trained mice performed a total of 500-1000 trials per daily session.

**Video monitoring and tracking body movements**

Mice were monitored during data acquisition using a Logitech HD Pro Webcam C920 (modified for IR detection), and a subset of the videos (15-30 fps) was used to track spontaneous body movements during the task with DeepLabCut[3]. Labels were manually placed at points of interest for tracking overall body motion (corner of the mouse stage), facial movements (snout tip, whisker pad and the two most prominently visible whiskers on either side of the face) and licking (tongue and tip of the lickport) to generate a training dataset. The deep neural network was trained for 200,000 iterations on 960 labeled frames extracted from 24 video clips across all three VIP-Cre; Ai162D mice. This training regimen produced a good fit of the model to the training data (loss <0.005). Behavioral videos were available for 12/23 imaging sessions included in analysis of VIP interneuron responses. These were used for analysis of VIP cell activity relative to whisk events.

Individual whisker identity was not tracked in the videos, rather an estimated of overall whisker motion over the session was quantified as the mean displacement of all whisker-related labels (whisker pad and two prominently visible whiskers on either side of the face), plotted as a function of time in the session. Whisk events were detected at timepoints when the whisker motion trace exceeded a threshold of the 85th percentile of displacement values for the entire session. Lick events estimated from the tongue and lickport labels were used to facilitate alignment of with behavior videos with trial timing information, relative to the pattern of licks detected by the capacitive lick sensor.

**2-photon calcium imaging**

Once criterion performance was achieved on the whisker detection task, imaging was conducted through the cranial window during daily behavior sessions. 2-photon imaging was performed using a Sutter Moveable-Objective Microscope with one resonant scanner (RESSCAN-MOM, Sutter) and one galvo scanner (Cambridge Technology). A Chameleon Ti-Sapphire pulsed laser (Coherent) tuned to 920 nm was used for excitation of GCamp6s. Fluorescence emission was collected through a water-dipping objective (16x, 0.8 NA, Nikon), band-pass filtered (HQ 575/50 filter, Chroma) and detected by GaAsP photomultiplier tubes (H10770PA-40, Hamamatsu). Single Z-plane images (512 x 512 pixels) were serially acquired at 7.5 Hz (30 Hz acquisition, averaged every 4 frames) using ScanImage 5 software (Vidrio Technologies). Laser power of 60-90 mW (at the front of objective) was used. Each imaging field measured 305 μm x 305 μm, and 7-8 imaging fields were obtained per mouse at depths of 110-250 μm below the dura, where the majority of VIP interneurons are found[4]. The blood vessels at the surface of each imaging field was imaged at the start of each session to for later use in alignment with vasculature during histology. After the final imaging session, the mouse was euthanized and the brain were collected for histology, to reconstruct the locations of imaging fields in the whisker map.

**Histology and localization of imaging fields**

The brain was removed and fixed overnight in 4% paraformaldehyde. The cortex was then flattened, sunk in 30% sucrose and sectioned at 50 µm parallel to the cortical surface. Tangential sections were processed for cytochrome oxidase (CO) staining. CO-stained sections revealed surface vasculature (in the most superficial section) and barrel boundaries (in sections through L4). Alignment of histological sections was performed manually using Fiji[5]. Custom MATLAB code was used to localize the centroid of each of the nine anatomical barrels that corresponding to whiskers stimulated per session, and the position of cells relative to barrel boundaries. Mean barrel width was calculated as the average of the major and minor axes across all barrels. Imaging fields were aligned to their locations in barrel cortex based on landmarks in surface vasculature, and the X-Y coordinate of each cell relative to barrel boundaries was determined. A cell was considered to be located within a specific barrel column if >50% of its pixels were located inside the boundaries of that column, as determined from the CO-stained sections. Cells not located within any column were classified as being located within septa. Distance of each cell from barrel column centroids was used for spatial analysis of sensory and action-related responses in the whisker map.

**Image processing and selection of ROIs**

Image processing was conducted in Matlab using custom pipeline code (adapted from LeMessurier, 2019[6]). Raw imaging movies were corrected for slow XY drift in Matlab using dftregistration[7]. Regions-of-interest (ROIs) were manually defined as ellipsoid regions drawn over neuronal somata that appeared when frames averaged across an entire registered imaging movie. The ROI signal was calculated as the mean fluorescence of its component pixels. Neuropil subtraction was not applied for VIP cells. For PYR cells included in supplemental analyses, neuropil correction was applied. A neuropil mask was created as a 10 pixel-wide ring starting two pixels from the somatic ROI. Neuropil pixels correlated with any somatic ROI (r>0.2) were excluded from the neuropil mask. Mean fluorescence of the neuropil mask was scaled by 0.3 and subtracted from the raw somatic ROI fluorescence. For each ROI, the raw fluorescence time series (or neuropil subtracted fluorescence trace, in the case of PYR cells) was converted to ΔF/F, which was defined as $(F_t-F_0)/F_0$, where $F_0$ is the $5^{th}$ percentile of fluorescence across the entire imaging movie and $F_t$ is the fluorescence on each frame.

**Analysis**

Behavioral performance:

Signal detection theory was applied to calculate *d'* as a measure of detection performance according to the standard definition as Z(Hit Rate) - Z(False Alarm Rate), where Z is the inverse cumulative of the normal distribution. For each session, analysis was restricted to trials between the first trial with a sliding d' > 0.5 to the last trial with a sliding d' > 0.5 to minimize effects of satiety level at the beginning and end of the testing session.

Quantification of evoked responses

Only sessions where a median of >25 trials per whisker were included for all analyses, to ensure accurate measurement of mean responses and receptive fields. All evoked responses were measured relative to the relevant event (stimulus, lick, whisking) using the same post-event analysis window (7 frames, 0.799 s) and pre-event baseline window (2 frames, 0.270 s).

On each Go trial, whisker-evoked ΔF/F signal was quantified for each ROI as (mean ΔF/F in the post-stimulus window) – (mean ΔF/F in 0 in the baseline window). On each NoGo trial, ΔF/F signal aligned to the NoGo stimulus (dummy piezo deflection) was calculated for each ROI, using the same method. For each ROI, the response magnitude to every whisker was calculated as (median whisker-evoked ΔF/F signal across Go trials – median ΔF/F signal across NoGo trials). Trials in which any licks occurred during the post-stimulus window were excluded from analysis. A cell was considered to have a significant whisker-evoked response if at least one whisker produced a significant response. A cell was considered whisker-responsive if it was significantly responsive to at least one whisker above baseline activity, as computed using a permutation test. To do this, the distribution of whisker-evoked ΔF/F signal on Go trials was combined with the distribution of median ΔF/F signal on NoGo trials. The combined distribution of mean signals was split into two groups, and the difference in means of the groups was measured. This was repeated 10,000 times, and the actual difference between the Go and NoGo distributions was compared to the distribution of permuted differences in means. A difference greater than the 95th percentile of the permuted distribution was considered to be significant. Correction for multiple comparisons (False Discovery Rate correction[8]) was applied to p-values for each of the nine whiskers.

To measure lick-evoked responses, the same procedure was followed, except that lick-evoked ΔF/F signal was used in place of whisker-evoked ΔF/F signals. The ΔF/F trace was aligned to lick times, as recorded by the capacitive lick sensor. Only the first lick of each lick bout were used (licks within bouts were excluded based on the distribution of interlick intervals on each trial). Licks were classified as ITI licks, FA licks or Hit licks depending on when they occurred in the trial structure. Only lick events that were not preceded by licks event for at least 3 s were included in the analysis of neural activity aligned to onset of whisk events. For ITI licks, an additional restriction of not being preceded by whisker stimulus for at least 3 s was also applied,

To measure responses evoked by spontaneous whisking in supplemental analyses, the same procedure used for stimulus- and lick-evoked responses was used, except that ΔF/F was aligned to whisk events determined from video analysis (described above in the section on 'Video monitoring and tracking body movements'). Only whisk events that were not preceded by whisk or licks event for at least 3 s were included in this analysis. Additionally, only whisk events that were not followed by a lick within 1 s were used – since the goal of this analysis was to examine activity evoked in VIP interneurons purely by whisking, as opposed to the whisker movements that typically occur while licking.

Quantification of receptive fields and map organization

The whisker associated with the anatomical home column of each cell was defined as its columnar whisker (CW). The whisker that evoked the numerically highest magnitude response was defined as the best whisker (BW). Equivalent best whiskers (eBWs) were defined as whiskers that evoked significant responses (relative to NoGo trials), but were not statistically different from the BW, based on permutation test. If the CW was amongst the eBWs for a cell, the cell was considered tuned to the CW, otherwise it was considered tuned to the BW. Organization of whisker-evoked vs. lick-related signals in the maps was examined as function of distance from the nearest column center. Cells in barrels and septa were both included in this analysis. Receptive field analyses were performed only for whisker-responsive cells. Negative ΔF/F responses were not excluded from receptive field analyses, however only cells with significant responses that had a positive BW response were included. Tuning heterogeneity within barrel columns was quantified as the fraction of cells within each column that were tuned to CW vs. non-CW whiskers. Septal cells were excluded from this analysis. To assess tuning heterogeneity across the whisker map, each cell's response to every whisker tested was examined as a function of distance from the center of the barrel column associated with that whisker (reference column). Whisker responses for each cell were normalized by Z-scoring to spontaneous (NoGo trial) activity, rank-ordered by response strength, and then averaged across cells to obtain the mean rank-ordered receptive field, centered on the BW. BW tuning preference was defined as $(R_{BW} - R_W)/( R_{BW} + R_W)$, where $R_{BW}$ = ΔF/F to BW, and $R_W$ = mean ΔF/F for all other whiskers. CW-centered tuning curves were also generated, as shown in supplementary analyses. CW tuning preference was defined as $(R_{CW} - R_W)/( R_{CW} + R_W)$, where $R_{CW}$ = ΔF/F to BW, and $R_W$ = mean ΔF/F for all other whiskers.

To verify that receptive field measures were not biased by the centering of the whisker array, all receptive analyses were also performed only for cells that had their CW positioned in the center of the piezo array (thus measuring responses to eight surrounding whiskers), however no significant differences were seen. Results are shown for the analysis of mean rank-ordered receptive field, using this method.

#### SUPPLEMENTAL FIGURE LEGENDS

**Figure S1.** Supplementary analyses of sensory and action-related responses of VIP interneurons in mouse whisker S1. **A.** Dynamics of VIP cell activity and licking behavior in the ITI period, prior to whisker stimulus onset (time 0). VIP cell activity (circles, n=3 mice) is compared with PYR cell activity (triangles) measured during the same task in one Drd3-Cre; Ai162D mouse. **B.** Mean ΔF/F trace for VIP cells aligned to spontaneous whisker movement (whisk) events compared to mean ΔF/F trace for the same cells aligned to lick events. Cells that were responsive to either whisker deflections or licks are included. **C.** Left, mean receptive fields for all L2/3 whisker-responsive VIP cells, separated into CW and ranked surround whiskers (SWs). Right, mean responses of VIP cells to CWs and SWs within the same row or the same arc.

### Figure S1

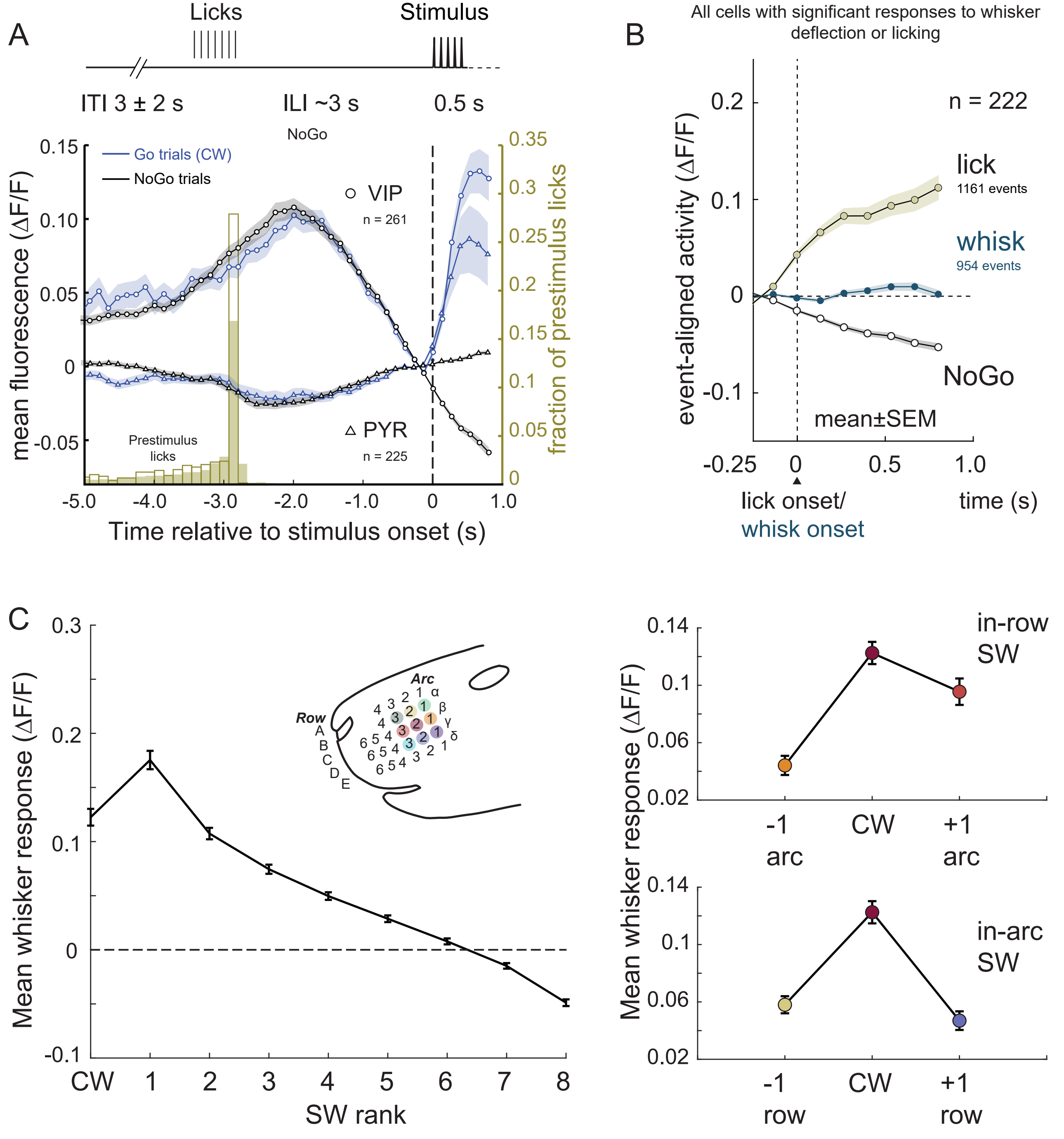
A
Licks
Stimulus
ITI 3 ± 2 s
ILI ~3 s
0.5 s
NoGo
Go trials (CW)
NoGo trials
VIP
n = 261
PYR
n = 225
Prestimulus
licks
mean fluorescence (ΔF/F)
fraction of prestimulus licks
Time relative to stimulus onset (s)
0.15
0.10
0.05
0
-0.05
0.35
0.3
0.25
0.2
0.15
0.1
0.05
0
-5.0
-4.0
-3.0
-2.0
-1.0
0
1.0
B
All cells with significant responses to whisker deflection or licking
n = 222
lick
1161 events
whisk
954 events
NoGo
mean±SEM
event-aligned activity (ΔF/F)
0.2
0.1
0
-0.1
-0.25
0
0.5
1.0
lick onset/
whisk onset
time (s)
C
Mean whisker response (ΔF/F)
0.3
0.2
0.1
0
-0.1
CW
1
2
3
4
5
6
7
8
SW rank
Arc
Row
A
B
C
D
E
1
2
3
4
5
6
α
β
γ
δ
in-row
SW
0.14
0.1
0.06
0.02
-1
arc
CW
+1
arc
in-arc
SW
-1
row
+1
row
